# Full-annual demography and seasonal cycles in a resident vertebrate

**DOI:** 10.1101/754929

**Authors:** Murilo Guimarães, Décio Correa, Marília Palumbo Gaiarsa, Marc Kéry

## Abstract

Demography is usually studied at a single point in time within a year when species, mostly long-distance migrants, are more active and easier to find. However, this provides only a low-resolution glimpse into demographic temporal patterns, compromising a complete understanding of species’ population dynamics over full annual cycles. The full annual cycle is often influenced by environmental seasonality, which induces a cyclic behavior in many species. However, cycles have rarely been explicitly included in models for demographic parameters, and most information on full annual cycle demography is restricted to migratory species. Here we used a high-resolution capture-recapture study of a resident tropical lizard to assess the full intra-annual demography and within-year periodicity in survival, temporary emigration and recapture probabilities. We found important variation over the annual cycle and up to 92% of the total monthly variation explained by cycles. Fine-scale demographic studies and assessments on the importance of cycles within parameters are fundamental to understand population persistence over time.

## Introduction

Demography is the study of four vital rates that underlie numerical change in populations: birth, death, immigration and emigration (Conroy and Carroll 2009). Understanding spatial and temporal patterns in these rates is essential for both ecology and management of living species; yet, we often lack sufficient resolution in our observations. For instance, studies on animal population dynamics usually focus on specific periods of the annual cycle, such as the breeding season, when a more sedentary lifestyle and increased activity enhance the probability of assessing individuals (Marra et al. 2015). This is the case for studies reporting individual variation in vital rates, such as survival probability, where individuals must be followed in the field over long periods (Lebreton et al. 1992). Although important, studies focusing on short periods typically can contribute less to our understanding of fundamental biological questions than do investigations extending over the full annual cycle of species (Marra et al. 2015 and references therein). Recently, Marra et al. (2015) have emphasized the importance of events beyond the breeding season and argued in favor of demographic research during the full annual cycle to fully understand specie’s population dynamics (Marra et al. 2015).

Most full-annual-cycle research focuses on a small handful of key periods within and outside of the breeding season (Hostetler et al. 2015), which for the vast majority of vertebrates, represents annual events. Although more expensive and time consuming than assessments made only during a single period within a year, models describing events over the full annual cycle are useful both for theoretical and applied questions including carry-over effects (Harrison et al. 2011), and detection of population trends (reviewed in Hostetler et al. 2015; Marra et al. 2015). Although most demographic models considering the full annual cycle have been applied to long-distance migratory species (Hostetler et al. 2015), especially birds (Culp et al. 2017; Rushing et al. 2017, but see Flockhart et al. 2015), species inhabiting the same area year-round offer better opportunities to study demography at a fine temporal scale. Nevertheless, resident species may also be subject to seasonal fluctuations due to less favorable periods for activity (Navas and Carvalho 2010), even in tropical areas (Dingle and Drake 2007).

Seasonal oscillations are expected for all living organisms (Panda et al. 2002) and are intrinsically connected to variability in temporal drivers, such as photoperiod (Bradshaw and Holzapfel 2007) or moon phase (Hauenschild 1960). For this reason, temporal predictors are commonly employed to explain individual activity and population demography. However, while temporal predictors are useful to understand the importance of temporal variability they may not always be available. Additionally, they may suffer from parameter estimability and are related to temporal variables, not to time itself (Fidino and Magle 2017). Within-year periodicity may often be fairly predictable, and accounting for such cyclicity in demographic rates may be better achieved by explicitly including cycles in population dynamics parameters. However, very rarely do population dynamics models include explicit within-season cycles and therefore, little is known about the magnitude of potential cyclic variation in population dynamics (Webster et al. 2002; Wingfield 2008; Hostetler et al. 2015; Marra et al. 2015). Our purpose here is to encourage research over the full annual cycle and to demonstrate periodicity may be intrinsic in vital parameters. We illustrate our point using a resident vertebrate, the neotropical whiptail lizard, *Ameivula ocellifera*, to investigate fine-scale, intra-annual demography through an intensive robust capture-recapture design (Pollock 1982; Kendall et al. 1997; Rankin et al. 2016). We first describe the temporal variation throughout the annual cycle and then, we evaluate the presence and magnitude of cyclic effects in demographic parameters of this resident neotropical species.

## Methods

### Study site and species

We conducted the study in SE Brazil, in a transitional area between the Cerrado and the Atlantic Forest biomes, in the Estação Ecológica de Jataí (21°37’3” S, 47° 45’55” W), state of São Paulo. Mean temperatures vary between 11 °C and 30 °C in the coldest and hottest months, respectively. Annual rainfall is around 1500 mm, mostly concentrated in the rainy season, from October to March (Cepagri 2011).

We surveyed a population of the whiptail lizard, *Ameivula ocellifera*, a small (30-78 mm SVL), and heliophilous Teiidae lizard that occurs in most parts of tropical Brazil (Mesquita and Colli 2003). Individuals are active year-round, especially in the rainy season (austral spring-summer) when breeding occurs. More details about the species may be found in Mesquita & Colli (2003).

### Sampling design

We sampled the population using a regular, square trapping grid consisting of 121 pitfall traps 25 m apart, over 6.2 ha, from September 2010 to September 2011, capturing individuals for seven consecutive days in each of 13 consecutive months. We used digital photography and batch marking, by clipping the third joint of the second toe of the right hand (ICMBio Animal Welfare Permit 10423-1), to recognize individuals. On each capture, we determined sex and assigned individuals to three sex/age classes: adult males (SVL > 40 mm), adult females (SVL > 51 mm), and newborns (SVL < 37 mm). After capturing and marking, we released individuals near the same trap where they had been caught. We used the Interactive Identification System software (Van Tienhoven et al. 2007) to identify individuals based on color and scale patterns. For more details, see Guimarães et al. (2017).

### Statistical Analysis

We modeled our mark-recapture data with the full-capture hierarchical robust design (RD) model (Rankin et al. 2016) and used a Bayesian mode of inference with MCMC techniques, to assess the effects of temporal variation on demographic parameters. The RD model distinguishes primary periods (here months), between which a population is assumed to be open, and nested secondary periods (here, daily capture occasions), for which a population is assumed to be closed.

We distinguished three demographic groups in our analyses: adult males, adult females, newborns. We expected different demographic rates for them and therefore, in our models, we stratified all parameters by these groups. In our analysis, we focused on three main biological parameters in the models: apparent survival probability (φ, hereafter ‘survival’), which is a product of true survival and site fidelity; temporary emigration probability, the probability of temporarily leaving the study site given the individual was onsite (γ”, hereafter ‘emigration’), and detection probability (*p*, hereafter, ‘recapture’), the probability of recapturing an individual given it is onsite. We also estimated abundances per group and month using parameter-expanded data augmentation (PX-DA, Royle and Dorazio 2012), which we do not focus on but describe in the Online Appendix 1. Our model also provides the probability of staying offsite (γ’), given the individual was offsite in the last sampling occasion and the probability of entering the study site (pent) along time. The former is hard to estimate and the latter is really more a nuisance parameter than of real biological interest (Rankin et al. 2016). Therefore, we do not discuss them in any detail.

We fit two variants of the RD model to our data that only differ in terms of the specification of the time variation in the parameters. In model 1, time was included as a random effect in parameters survival, emigration and recapture. That is, in this model we allowed parameters to be different for each month and to vary around a constant value according to a random effect. In model 2, we added cyclic effects on the same three parameters, survival, emigration, and recapture, using a periodic cosine function of month, and on top of that specified monthly residuals, also as random effect. Our approach is similar to Flury and Levri (1999), where we added two effects of month into the linear predictor of these three parameters, as follows: 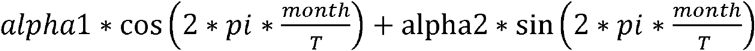 where *pi* is the number Pi, *month* corresponds to the month number between 1 (January) and 12 (December), *T* represents the length of the periodic response (here *T*=12), and *alpha1* and *alpha2* are the two coefficients associated with the covariate ‘month’. In this formulation, *month/T* tells us how far through a full annual cycle of 2*pi each month is (see model code in the Online Appendix 2). In model 2, the proportional reduction in the among-month variation achieved by specification of the cyclic patterns in these parameters enabled us to quantify the proportion of the among-month variation that could be explained by the cycles (Kéry and Schaub 2012).

We used vague priors and hyperpriors for all parameters, which we show in the Online Appendix 2. We implemented our models in the BUGS language (Lunn et al. 2000) using program JAGS (Plummer 2003), run from R (R Core Team 2018) through the jagsUI interface (Kellner 2014). We ran three chains with 400,000 iterations each, discarding 300,000 as burn-in and thinning at a rate of 1 in 100. We determined chain convergence by visual examination of trace plots and by the Brooks-Gelman-Rubin statistic (Brooks and Gelman 1998), which was < 1.1 for all parameters. We present posterior means and 95% credible intervals (CRI).

## Results

Our field effort consisted of 11,011 trap-days during which we captured 164 adult males, 163 adult females and 89 newborns for a total of 416 individuals. Fifty-one adult males, 29 adult females and 14 newborns were recaptured at least once.

Under model 1 (with random month effects), mean monthly apparent survival probability was much higher for adult males (0.94, CRI 0.81-0.99) and adult females (0.90, CRI 0.72-0.99) than for newborns (0.64, CRI 0.17-0.96). We observed considerable variability in survival during the course of a year, where adults of both sexes presented similar patterns through the year, and newborns showed a sharp decrease after hatchling (Fig. 1).

**Fig. 1.**
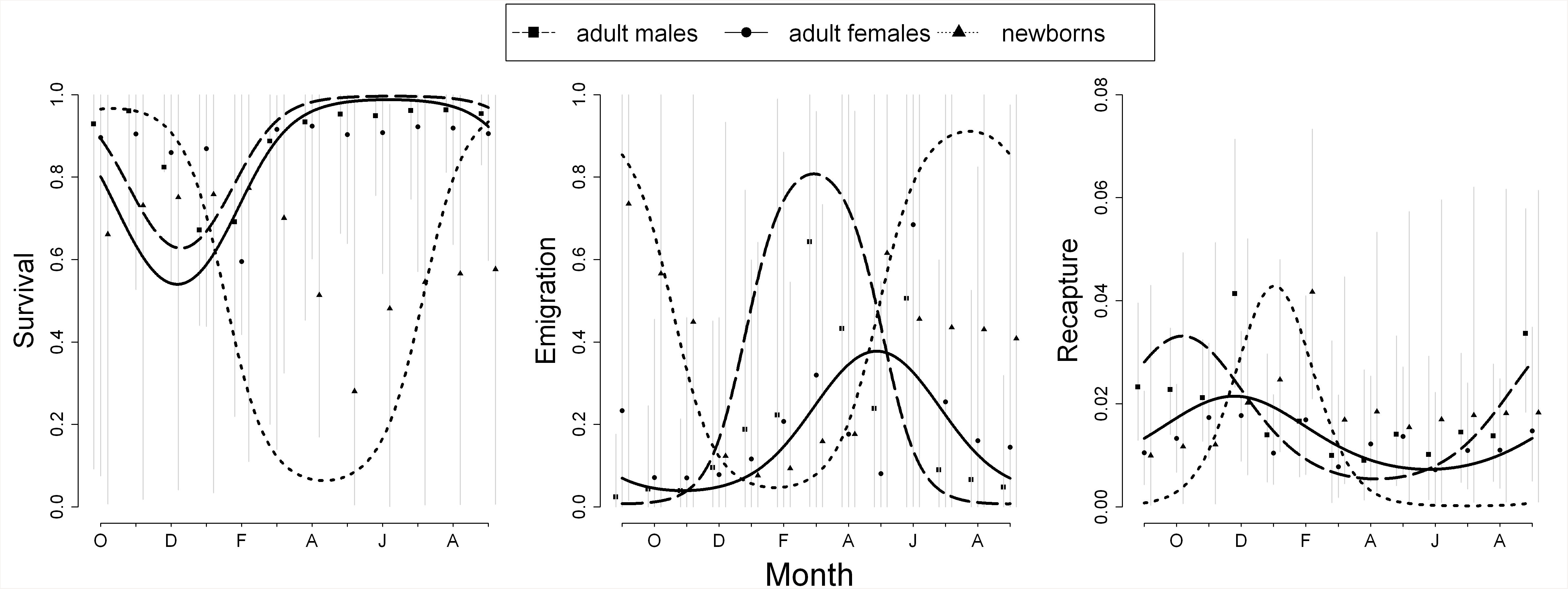
Probability of survival, emigration and recapture for adult males, adult females and newborns for each month based on model 1 (symbols and gray credible interval vertical lines, random month effects only) and model 2 (solid and dashed lines, cycles plus random monthly deviations from these cycles [see text for details]).

The newborns showed the highest probability of emigration (0.32, CRI 0.01-0.85), followed by adult females (0.16, CRI 0.03-0.63) and males (0.15, CRI 0.042-0.66, Fig. 1). Daily mean recapture probabilities were similar in all three groups with small variations during the year, where males and newborns presented the highest rates during the breeding season, at the beginning and in the end, respectively (Fig. 1).

Model 2 (with cycles added) revealed a strong periodicity in all three parameters that we modeled with a cosine periodic function of the month (survival, emigration and recapture; Fig. 1). Comparisons of the magnitude of the (residual) among-month variance between models 2 and 1 showed that annual cycles explained 92.5% of the monthly variation in apparent survival, 86.8% in emigration probability, and 84.6% in recapture probability.

## Discussion

Most population dynamics studies are based on the observation of a population just once a year, often during the breeding season. Recently, it has been pointed out that studies with high temporal resolution are required to fully understand population dynamics, so are models covering the full annual cycle of a species, i.e., with multiple samplings a year (Hostetler et al. 2015; Marra et al. 2015). Here, we modeled fine-scale temporal variability in the vital rates of a resident species and assessed the population throughout the year, including low-activity periods. We showed that demographic parameters varied along the full annual cycle. Furthermore, we detected a strong periodic signal in those vital parameters demonstrating cyclicity is intrinsically associated to demography. Our findings broaden the understanding of specie’s population biology and were only possible due to the unusually high-temporal resolution of demographic information for an entire annual cycle.

### The full annual cycle reveals detailed information on species biology

Survival estimates for the vast majority of the ~ 6,500 species of lizards are lacking, but general patterns have emerged based on life history traits, including foraging mode and mating system. Our results contribute to the relationship between clutch size and growth rate – the classical slow-fast continuum –, which finds support in lizard natural history where early investments in rapid growth may decrease late survival (Clobert et al. 1998; Olsson and Shine 2002). Survival in slow-growing, territorial lizards from the *Iguania* clade are said to be higher (Ujvari et al. 2015; Iverson et al. 2016; Keehn et al. 2019), than in the non-territorial *Scleroglossa* clade, to which the *Teiidae* family belongs (Pianka and Vitt 2003). Males from Teiidae species can present female accompaniment (Pianka and Vitt 2003) and fight conspecific males to guard females, which may impair survival (Ancona et al. 2010). Females benefit from mate guarding, but may present survival decrease due to basking activity and low mobility during pregnancy (Shine 1980). We found that survival dropped 30% to 35% after the breeding season for adult males and females, respectively. Thus, our results support the hypothesis that reproduction is costly and negatively affects future survival, further highlighting the importance of full annual cycle research.

Temporary emigration for adults was the highest after the mating and hatchling period, when energetic reserves may be depleted (Shine 1980). This period of the full-annual cycle also characterizes the beginning of the dry and cold season, when we found several individuals buried 10-20 cm below ground. Newborn’s temporary emigration probability was up to four times higher than that of adults explaining, at least in part, the scarcity of juvenile demographic information that is common across taxonomic groups. This pattern is usually attributed to different aspects, including smaller body sizes, secretive and inconspicuous habits, and high mortality rates (Rivas et al. 2016; Bailey et al. 2017; Wilson et al. 2018).

Adult males presented the highest recapture rates at the onset of the breeding season, which coincides with high mating activity rates during this period. Since our trap system was stationary, we would expect capture rates to increase with activity, and in this case, detection probability parameter of a capture-recapture model is not merely a nuisance parameter. More than that, it is a parameter with a biological meaning, related to physical activity, analogous to singing activity, which produces a strong signal in the detection probability of passerine birds (Strebel et al. 2014). In territorial species, males may present higher recapture rates than females due to greater site fidelity. Overall, our mean recapture estimates were similar across groups. Given that this species feeds primarily on termites, where all individuals actively forage in open areas (Mesquita and Colli 2003), our results may be related to their feeding behavior and the absence of territoriality (Pianka and Vitt 2003).

### Periodicity can be strong within demographic rates

Cyclicity is found in endogenous rhythms of every life form, from cells to individuals, and the biological clock is influenced by exogenous rhythms, such as Earth’s rotation and climate, to maximize fitness (Panda et al. 2002; Wingfield 2008). This explains, at least in part, why activity in many organisms is aggregated in short time frames, such as breeding events (Bradshaw and Holzapfel 2007; Harrison et al. 2011; Marra et al. 2015). Although the full annual cycle is mostly described for long-distance migratory species, where individuals are unobservable for some time, tropical species may also be subject to periodicity due to temporary absences, involving reduced activity and unavailability.

Surprisingly, few studies exploring species’ life cycle have actually fitted cyclic patterns in specie’s demographic rates. If we assume that the beginning of a temporal pattern in a parameter to “meet” with the end of it over the course of a year, such cycles are almost to be expected. Periodicity may occur at different time scales, such as observed in the daily activity of the New Zeland mudsnails (Flury and Levri 1999), or in the yearly activity of American mesocarnivore mammals, with up to 75% of temporal variability in colonization rates explained by ciclicity (Fidino and Magle 2017). We fitted cycles in three demographic parameters (apparent survival, emigration probability and recapture rate) and found strong evidence of annual periodicity, with cycles explaining on average 88% of the total variation among months. Even though the observed variability in the demographic parameters was based on only one full cycle, we claim they can describe monthly variations within a year and may be of great use to wildlife management and conservation. Further information on population demography may help support our findings on the magnitude of specie’s annual cycles.

In the face of rapid biodiversity declines, information on individuals and populations are essential for conservation purposes. Full annual cyclic modeling, the combination of full annual cycle research with cyclic parameter estimation, can be powerful to uncover sensitive points in the trajectory of populations, supporting decision-making actions. Our results illustrate two important points; first, we endorse and encourage the use of full annual cycle studies of species, emphasizing investigations on resident species, since they can provide cheaper fine-scale information than long-distant migratory species. Second, we demonstrated that intra-annual temporal variability in demographic rates is highly cyclic. Periodicity has rarely been explicit included in vital rate’s studies, and here we claim for its inclusion in population demography as a venue for predictive studies. Fluctuations may be triggered by many different sources, and understanding the mechanisms that regulate populations over time is of fundamental importance to ensure specie’s persistence. Full annual cyclic modeling may be important to predict future population oscillations and raise important insights, even when they explain little variability.

## Acknowledgements

We thank Robert Rankin and Brett McClintock for advice on statistical analysis. Ricardo Sawaya, Hamanda B. Cavalheri, Sérgio Serrano, Thiago Oliveira, Roberto Munguia Steyer, Glauco Machado and our field crew were especially kind and helpful during data collection. We thank ICMBio for the permission on data collection and CNPq (140684/20093), CAPES (2296/110), FAPESP (2008/54472-2), the Ecology Graduate Program of UNICAMP and the Schifferli Scholarship Program of the Swiss Ornithological Institute for financial support.

## References

Ancona S, Drummond H, Zaldívar-Rae, J (2010) Male whiptail lizards adjust energetically costly mate guarding to male-male competition and female reproductive value. Anim Behav 79: 75–82

Bailey AM, McCleery RA, Ober HK, Pine WE (2017) First demographic estimates for endangered Florida bonneted bats suggest year-round recruitment and low apparent survival. J Mammal 9:551–559

Bradshaw WE, Holzapfel CM (2007) Evolution of animal photoperiodism. Annu Rev Ecol Evol Syst 38:1–25

Brooks SP, Gelman A (1998) General methods for monitoring convergence of iterative simulations. J Comput Graph Stat 7:434–455

Clobert J, Garland JrT, Barbault R (1998) The evolution of demographic tactics in lizards: a test of some hypotheses concerning life history evolution. J Evol Biol 11:329–364

Conroy MJ, Carroll JP (2009) Quantitative Conservation of Vertebrates. (MJ Conroy, JP Carroll, Eds.) (First). Oxford: Wiley-Blackwell

Culp LA, Cohen EB, Scarpignato AL, Thogmartin WE, Marra PP (2017) Full annual cycle climate change vulnerability assessment for migratory birds. Ecosphere, 8: 1–22

Dingle H, Drake VA (2007) What is Migration? Can Stud Popul 57:113–121

Fidino M, Magle SB (2017) Using Fourier series to estimate periodic patterns in dynamic occupancy models. Ecosphere 8:9

Flockhart DTT, Pichancourt JB, Norris DR, Martin TG (2015) Unravelling the annual cycle in a migratory animal: breeding-season habitat loss drives population declines of monarch butterflies. J Anim Ecol 84:155–165

Flury BD, Levri EP (1999) Periodic logistic regression. Ecology 80:2254–2260

Guimarães M, Munguía-Steyer R, Doherty PF, Sawaya RJ (2017) No survival costs for sexually selected traits in a polygynous non-territorial lizard. Biol J Linn Soc 122:614–626

Harrison XA, Blount JD, Inger R, Norris DR, Bearhop S (2011) Carry-over effects as drivers of fitness differences in animals. J Anim Ecol 80:4–18

Hostetler JA, Sillett TS, Marra PP (2015) Full-annual-cycle population models for migratory birds. Auk 132:433–449

Iverson JB, Smith GR, Pasachnik SA, Hines KN, Piepper L (2016) Growth, coloration, and demography of an introduced population of the Acklinns Rock Iguana (Cyclura rileyi nuchalis) in the Exuma Islands, The Bahamas. Herpetol Cons Biol 11:139–153

Keehn JE, Shoemaker KT, Feldman CR (2019) Population-level effects of wind farms on a desert lizard. J Wild Manag 83:145–157

Kendall WL, Nichols JD, Hines JE, Mar N (1997) Estimating temporary emigration using capture-recapture data with Pollock’s Robust Design. Ecology, 78:563–578

Kéry M, Schaub M (2012) Bayesian population analysis using WinBUGS. (MKéry, MSchaub, Eds.) (First). Amsterdam: Elsevier

Lebreton J-D, Burnham KP, Clobert J, Anderson DR (1992) Modeling survival and testing biological hypotheses using marked animals: a unified approach with case studies. Ecol Monogr 62:67–118

Lunn DJ, Thomas A, Best N, Spiegelhalter D (2000) WinBUGS - a bayesian modelling framework: concepts, structure, and extensibility. Stat Comput 10:325–337

Marra PP, Cohen EB, Loss SR, Rutter JE, Tonra CM (2015) A call for full annual cycle research in animal ecology. Biol Lett 11:20150552

Mesquita DO, Colli GR (2003) The Ecology of Cnemidophorus ocellifer (Squamata, Teiidae) in a Neotropical Savanna. J Herpetol 37:498–509

Navas CA, Carvalho JE (2010) Aestivation: molecular and physiological aspects. (CA Navas, JE Carvalho, Eds.) (First). Heildelberg: Springer

Olsson M, Shine R (2002) Growth to death in lizards. Evolution, 56:1867–1870.

Panda S, Hogenesch JB, Kay SA (2002) Circadian rhythms from flies to human. Nature, 417:329–335

Pianka E, Vitt L (2003) Lizards: windows to the evolution of diversity. (E Pianka, LJ Vitt, Eds.). Berkeley: University California Press

Plummer M (2003) JAGS: a program for analysis of Bayesian graphical models using Gibbs sampling. In K. Hornik, F. Leisch, & A. Zeileis (Eds.), 3rd International Workshop on Distributed Statistical Computing 3:1–10

Pollock KH (1982) A capture-recapture sampling design robust to unequeal catchability. J Wild Manag 46:752–757

Rankin RW, Nicholson KE, Allen SJ, Krützen M, Bejder L, Pollock KH (2016) A Full-Capture Hierarchical Bayesian Model of Pollock’s Closed Robust Design and Application to Dolphins. Front Mar Sci 3:1–18

Rivas JA, Molina CR, Corey SJ, Burghardt GM (2016) Natural history of neonatal green anacondas (Eunectes murinus): a chip off the old block. Copeia, 104:402–410

Royle JA, Dorazio RM (2012) Parameter-expanded data augmentation for Bayesian analysis of capture-recapture models. J Ornithol 152:521–537

Rushing CS, Hostetler JA, Sillett TS, Marra PP, Rotenberg JA, Ryder TB (2017) Spatial and temporal drivers of avian population dynamics across the annual cycle. Ecology 98:2837–2850

Shine R (1980) “Costs” of reproduction in reptiles. Oecologia, 46:92–100

Strebel N, Kéry M, Schaub M, Schmid H (2014) Studying phenology by flexible modelling of seasonal detectability peaks. Methods Ecol Evol 5:483–490

Ujvari B, Fisher P, Rydell J, Wahlgren R, Wright B, Madsen T (2015) Population demography of frillneck lizards (Chlamydosaurus kingii, Gray 1825) in the wet-dry tropics of Australia. Austral Ecol 40:60–66

Van Tienhoven AM, Den Hartog JE, Reijns RA, Peddemors VM (2007) A computer-aided program for pattern-matching of natural marks on the spotted raggedtooth shark Carcharias taurus. J Appl Ecol 44:273–280

Webster MS, Marra PP, Haig SM, Bensch S, Holmes RT (2002) Links between worlds: unraveling migratory connectivity. Trends Ecol Evol 17:1–8

Wilson S, Saracco JF, Krikun R, Flockhart DTT, Godwin CM, Foster KR (2018) Drivers of demographic decline across the annual cycle of a threatened migratory bird. Sci Rep 8:1–11

Wingfield JC (2008) Organization of vertebrate annual cycles: implications for control mechanisms. Philos Trans R Soc Biol Sci 363:425–441

