## Supplemental material for "Full-annual demography and seasonal cycles in a resident vertebrate"

**Electronic Supplemental Material**

### Supplemental material 1: abundance estimation procedure

We estimated abundance, using the parameter-expanded data augmentation (PX-DA, Royle and Dorazio 2012), where we added an arbitrary but large number of all-zero capture histories to the data (Royle and Dorazio 2012), bringing the total number of individuals in the augmented data set to size $M$. The $M – n$ augmented individuals (where $n$ is the observed number of individuals), represent *potential* individuals that could be alive and within the study area. The model allows us to estimate the proportion of individuals that are actually associated with the study area. Another way to describe PX-DA in the Bayesian context is simply to note that this is a manner of specifying a discrete uniform prior between 0 and $M$ for population size N (Chapter 6 in Kéry and Schaub 2012). To make sure we augmented enough pseudo-individuals, we checked the cumulative probability of inclusion, making sure its upper 95% credible interval was always below 0.7 (R. Rankin, pers. com.). Finally, we estimated recruitment probability ($\psi$), representing the transition from "not yet in the population" to "alive and at risk of capture". In this context, the recruitment rate includes an unobservable state and is challenging to estimate in the context of our model (Rankin et al. 2016), because the way new recruits enter the population (on either offsite or onsite states) has to be specified. Here, we set new recruits to have the same movement behavior as individuals already captured and marked in the population (Rankin et al. 2016). Due to technical reasons (Rankin et al. 2016), we constrained one of the parameters, $\gamma$', to be constant over time, while we allowed full time-dependence for all other parameters to enable us to investigate temporal patterns in them.

Population size peaked during the first months of the breeding season (end of austral winter) when it reached 590 (CRI 390-890) individuals and was lowest in the dry/cold period, with 170 (CRI 27-400) individuals. The considerable uncertainty in the latter estimate is due to the low detection probability in that season. Females were more abundant than were males and newborns during the whole year. The number of newborns peaked in the end of hatchling period and approached zero close to the next mating season (Fig. S1).

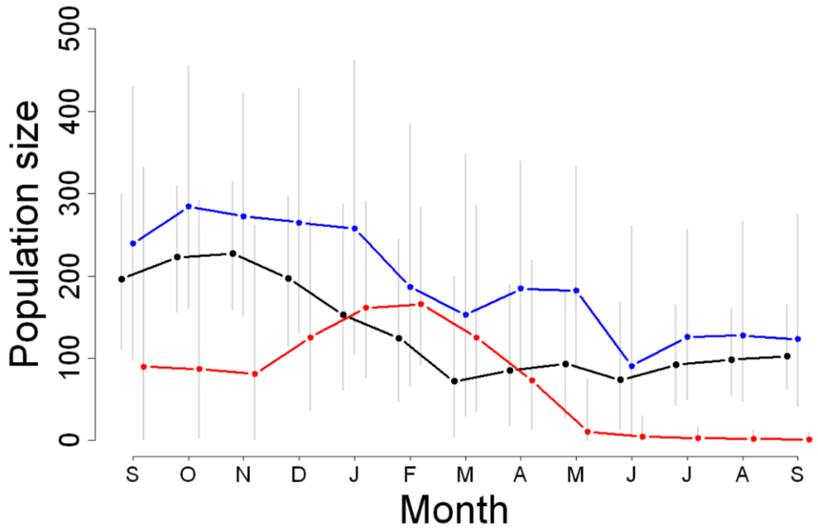


**Fig. S1** Abundance estimates for adult males (black), adult females (blue) and newborns (red) based on model 1 (random month effects only). Gray lines indicate 95% credible intervals.

### Supplemental material 2: R code and JAGS model

```
# This code replicates the analyses of mark-recapture data using the Robust Design adapted from Rankin
# et al. (2017) to BUGS language. Please, read Rankin et al. (2017) before running the analysis.
# This code is focused on the estimation of survival, emigration and recapture probabilities
# using a cycle predictor. Code was adapted by Marc Kéry and Murilo Guimarães


library(jagsUI)

#Load files
source("R_PCRD_JAGS_SOURCE.R") # load handy functions from Rankin et al.(2017) paper
MARK.file.name <- "RD_whiptail.inp" #load inp file

#Bundle and summarize data set
str(bdata <- list(y=Ysum, T=T, M=mm, group = group, pi = pi, month.phi = 0:11, month.p.gamma = 0:12))
#List of 7
# $ y            : num [1:1664, 1:13] 1 1 1 1 1 1 1 1 1 1 ...
#  ..- attr(*, "dimnames")=List of 2
#  .. ..$ : chr [1:1664] "M1" "M2" "M3" "M4" ...
#  .. ..$ : NULL
# $ T            : int 13
# $ M            : int 1664
# $ group        : num [1:1664] 1 1 1 1 1 1 1 1 1 1 ...
# $ pi           : num 3.14
# $ month.phi    : int [1:12] 0 1 2 3 4 5 6 7 8 9 ...
# $ month.p.gamma: int [1:13] 0 1 2 3 4 5 6 7 8 9 ..


# Write model in BUSG language
sink("model5.txt")
cat("
    model{
    ### Priors and linear models
    # ----------------------------------------------
    # Group- and time-dependent survival (interaction effects)
    # First survival is redundant and will be sampled from U(0,1)
    for(g in 1:3){
    phi[g,1] ~ dunif(0, 1)
    } # g
    # Survival for intervals 2:13 are 'real'
    for(t in 2:T){ # Loop over T=13 primary periods...
    for(g in 1:3){
    phi[g,t] <- ilogit(lphi[g,t])     #apparent survival probability on logit scale
    lphi[g,t] ~ dnorm(mu.lphi.group[g] + beta1.phi[g] * cos(2*pi*month.phi[t-1] / 12) + beta2.phi[g] *
     sin(2*pi*month.phi[t-1] / 12) , tau.lphi.time)     # Different mean, but same variance
    phi_cycle[g,t-1] <- ilogit(mu.lphi.group[g] + beta1.phi[g] * cos(2*pi*month.phi[t-1] / 12) +
     beta2.phi[g] * sin(2*pi*month.phi[t-1] / 12))
    } # g
    } # t
    for(g in 1:3){
    mu.lphi.group[g] <- logit(mean.phi[g])
    mean.phi[g] ~ dunif(0, 1)
    beta1.phi[g] ~ dnorm(0, 0.1)
    beta2.phi[g] ~ dnorm(0, 0.1)
    }
    tau.lphi.time <- pow(sd.lphi.time, -2)
    sd.lphi.time ~ dt(0, 0.1, 5)I(0, )       # Half-t prior for variance (in sd scale)

    # Group-dependent gamma1 constant over time
    pr.gamma1 <- c(1.3,1.3)
    for(g in 1:3){
    gamma1[g] ~ dbeta(pr.gamma1[1],pr.gamma1[2])
    } # g
    # Group and time-dependent gamma2 (with interaction, and with cycles as well now)
    for(t in 1:T){ # Loop over T=13 primary periods
```

```
for(g in 1:3){   # Loop over 3 groups
gamma2[g,t] <- ilogit(lgamma2[g,t]) # temp migration
lgamma2[g,t] ~ dnorm(mu.lgamma2.group[g] + beta1.gamma2[g] * cos(2*pi*month.p.gamma[t] / 12) +
 beta2.gamma2[g] * sin(2*pi*month.p.gamma[t] / 12) , tau.lgamma2.time)
gamma2_cycle[g,t] <- ilogit(mu.lgamma2.group[g] + beta1.gamma2[g] * cos(2*pi*month.p.gamma[t] / 12)
 + beta2.gamma2[g] * sin(2*pi*month.p.gamma[t] / 12))
} # g
} # t
for(g in 1:3){
mu.lgamma2.group[g] <- logit(mean.gamma2[g])
mean.gamma2[g] ~ dunif(0, 1)
beta1.gamma2[g] ~ dnorm(0, 0.1)
beta2.gamma2[g] ~ dnorm(0, 0.1)
}
tau.lgamma2.time <- pow(sd.lgamma2.time, -2)
sd.lgamma2.time ~ dt(0, 0.1, 5)I(0, )   # Half-t prior for variance (in sd scale)
# Prior for vector of probabilities for the categorical to estimate group membership
for (g in 1:3){
beta[g] ~ dgamma(1, 1)   # Induce Dirichlet prior
theta[g] <- beta[g] / sum(beta[])
} # g
# Group and time-dependent detection:  (interaction effects, with cycles)
for(t in 1:T){ # Loop over T=13 primary periods...
for(g in 1:3){   # Loop over 3 groups
p[g, t] <- ilogit(lp[g, t])     # Define logit of detection probability
lp[g,t] ~ dnorm(mu.lp.group[g] + beta1.p[g] * cos(2*pi*month.p.gamma[t] / 12) + beta2.p[g] *
 sin(2*pi*month.p.gamma[t] / 12) , tau.lp.time)
p_cycle[g,t] <- ilogit(mu.lp.group[g] + beta1.p[g] * cos(2*pi*month.p.gamma[t] / 12) + beta2.p[g] *
 sin(2*pi*month.p.gamma[t] / 12))
} # g
} # t
for(g in 1:3){
mu.lp.group[g] <- logit(mean.p[g])
mean.p[g] ~ dunif(0, 1)
beta1.p[g] ~ dnorm(0, 0.1)
beta2.p[g] ~ dnorm(0, 0.1)
}
tau.lp.time <- pow(sd.lp.time, -2)
sd.lp.time ~ dt(0, 0.1, 5)I(0, )    # Half-t prior for variance (in sd scale)
# ------------- Model for group membership -------------------------------
for(i in 1:M){
group[i] ~ dcat(theta[1:3])
}
# Model for recruitment process from eigenvector decomposition
for(t in 1:T){
psi[t] ~ dunif(0,1)   # inclusion probability
}
for(t in 1:T){
for(i in 1:M){
lambda[1,i,t] <- (1-gamma1[group[i]])/(gamma2[group[i],t]- gamma1[group[i]] +1)
lambda[2,i,t] <- 1-lambda[1,i,t] # long-term prob of being outside
# trmat: transition matrix for Markovian latent-states
# 1 =not yet in population;2=dead;3=offsite;4=onsite (only observable state)
# trmat[row,column,time] = [state at time=t (arrival); state at time t-1 (departure); i #=
 individual, time=t]
trmat[1,1,i,t] <- 1-psi[t]                 # excluded from pop
trmat[2,1,i,t] <- 0                        # dead
trmat[3,1,i,t] <- psi[t]*lambda[2,i,t]     # inclusion into pop, outside study are
trmat[4,1,i,t] <- psi[t]*lambda[1,i,t]     # inclusion into pop, inside study area
trmat[1,2,i,t]<- 0
trmat[2,2,i,t]<- 1                         # stay dead
trmat[3,2,i,t]<- 0
trmat[4,2,i,t]<- 0
trmat[1,3,i,t]<- 0
trmat[2,3,i,t]<- 1-phi[group[i],t]                      # dies outside
trmat[3,3,i,t]<- gamma1[group[i]]*phi[group[i],t]       # stays outside | outside
trmat[4,3,i,t]<- (1-gamma1[group[i]])*phi[group[i],t]   # reenters study area | outside
trmat[1,4,i,t]<- 0                                      # stays dead
trmat[2,4,i,t]<- 1-phi[group[i],t]                      # dies inside
trmat[3,4,i,t]<- gamma2[group[i],t]*phi[group[i],t]     # leaves study area | inside
trmat[4,4,i,t]<- (1 - gamma2[group[i],t])*phi[group[i],t]  # stays inside | inside
} #i
} # t
# likelihood: loop through M individuals, both real and pseudo-individuals
```

```
    for (i in 1:M){
    #draw latent state at primary period 1:
    # ... by definition, everyone starts in z=1 (not-in-population) at time=0
    z[i,1] ~ dcat(trmat[1:4,1,i,1]) # first z strictly excluded from pop
    # likelihood for first primary period
    # Binomial observation process, conditional on z=4, otherwise no observation
    y[i,1] ~ dbinom(p[group[i], 1] * equals(z[i,1], 4), 7)
    # loop through primary periods after 1st primary periods
    for(t in 2:T){
    # state process: draw z(t) conditional on z(t-1)
    z[i,t] ~ dcat(trmat[1:4, z[i,t-1] , i, t])
    # likelihood
    # Binomial observation process, conditional on z=4, otherwise no observation
    y[i,t] ~ dbinom(p[group[i], t] * equals(z[i,t],4), 7)
    } # t
    } # i
    # estimate population size per primary periods
    for(t in 1:T){
    for (i in 1:M){
    # tally up alive individuals (in states 3 and 4) and those that are inside of the study area (i.e.,
   state 4)
    alive_i[i,t] <- step(z[i,t]-3) # check alive or not
    Nin_i[i,t] <- equals(z[i,t],4) # count if i is within study area
    Nin_i1[i,t] <- equals(z[i,t],4) * equals(group[i],1) # count if i is within study area
    Nin_i2[i,t] <- equals(z[i,t],4) * equals(group[i],2) # count if i is within study area
    Nin_i3[i,t] <- equals(z[i,t],4) * equals(group[i],3) # count if i is within study area
    } # i
    alive[t] <- sum(alive_i[,t]) # number alive
    Nin[t] <- sum(Nin_i[,t]) # number in study area
    Nin1[t] <- sum(Nin_i1[,t]) # number in study area and in group 1
    Nin2[t] <- sum(Nin_i2[,t]) # number in study area and in group 2
    Nin3[t] <- sum(Nin_i3[,t]) # number in study area and in group 3
    } # t
    # Calculate POPAN pent (probability of entry)
    cumprob[1] <- psi[1]
    for(t in 2:T){
    cumprob[t] <- psi[t]*prod(1-psi[1:(t-1)])
    }
    cumprob.norm <- sum(cumprob[1:T])
    # POPAN probabilities
    for(t in 1:T){
    pent[t] <- cumprob[t]/cumprob.norm
    } #t
    }", fill=TRUE)
sink()
```
